# Identification of subgroups and development of prognostic risk models along the glycolysis-cholesterol synthesis axis in lung adenocarcinoma

**DOI:** 10.1101/2022.08.07.503114

**Authors:** Bao Qian, Yangjie Guo, Jiuzhou Jiang

## Abstract

**Background:** Lung cancer is one of the most dangerous malignant tumors affecting human health. Lung adenocarcinoma (LUAD) is the most common subtype of lung cancer. Both glycolytic and cholesterogenic pathways play critical roles in metabolic adaptation to cancer.

**Methods:** A dataset of 585 LUAD samples was downloaded from The Cancer Genome Atlas (TCGA) database. We obtained co-expressed glycolysis and cholesterogenesis genes by selecting and clustering genes from Molecular Signatures Database v7.5. Compared the prognosis of different subtypes and identified differentially expressed genes (DEGs) between subtypes. Predictive outcome events were modeled using machine learning, and the top 9 most important prognostic genes were selected by Shapley additive explanation (SHAP) analysis. A risk score model was built based on multivariate Cox analysis.

**Results:** LUAD patients were categorized into four metabolic subgroups: cholesterogenic, glycolytic, quiescent, and mixed. The worst prognosis was the mixed subtype. The prognostic model had great predictive performance in the test set.

**Conclusions:** Patients with LUAD were effectively typed by glycolytic and cholesterogenic genes and were identified as having the worst prognosis in the glycolytic and cholesterogenic enriched gene groups. The prognostic model can provide an essential basis for clinicians to predict clinical outcomes for patients.

## 1. Introduction

Although lung cancer is the second most common cancer globally, it is the leading cause of cancer deaths, accounting for an annual estimated total of two million new cases and 1.76 million deaths [1,2]. Lung cancer can be broadly grouped into small-cell lung cancer (SCLC, 15%) and non-small-cell lung cancer (NSCLC, 85%), and lung adenocarcinoma (LUAD) is the most common subtype of lung cancer [2–4]. The treatment of NSCLC has changed dramatically over the past decade, primarily due to advances in biomarkers that allow for targeted and immune-based therapies for specific patients with significant success [5]. However, the vast majority of advanced NSCLC become resistant to current treatments and eventually progress [6]. Therefore, searching for new predictors to predict and improve the prognosis of LUAD is imminent.

Reprogramming of cell metabolism is an essential feature of malignancy, as shown by abnormal uptake of glucose and amino acids and dysregulation of glycolysis [7,8]. Glycolysis is a specific metabolic pattern of tumor cells, which meets the requirements of tumor cells for ATP, etc.[9]

The analysis of large volumes of complex biomedical data through computer algorithms, driven by the ongoing development of computer hardware and enormous amounts of data, offers substantial advantages for advancing biology and accurately estimating patient conditions [10,11]. Machine learning (ML) is a scientific discipline focusing on how computers learn from data and build predictive models. It is becoming an embedded part of modern research in biology, but its “black box” nature is an additional challenge. [12,13]. Interpretation is an integral branch of method development, with Shapley additive explanation (SHAP) being an integral approach [14]. The SHAP, which explains the model outcome by computing the contribution of each input feature for all samples, was applied to study the effects of different variables[15,16]. A positively or negatively valued SHAP represents an increase or decrease in the probability of a specific outcome.

Balanced glycolysis and cholesterol production pathways can jointly regulate tumor progression. However, there is a lack of studies to establish LUAD-related staging based on the glycolysis-cholesterol synthesis axis. So based on the gene expression levels of glycolysis-cholesterogenesis, we defined four subtypes of LUAD from a metabolic perspective and further analyzed the characteristics of different subtypes, such as survival time and other clinical features in this study. Then, we identified prognostic genes based on machine learning, further explored the correlation of the genes with clinical features and tumor microenvironment (TME) and proposed a risk-prognosis model that can be used to formulate treatment options and analyze the prognosis.

## 2. Materials and Methods

### Data acquisition and processing

Gene expression data (count matrix), corresponding clinical information, and the somatic mutation profiles for 585 LUAD patients were downloaded from The Cancer Genome Atlas (TCGA) via the UCSC Xena browser (https://xenabrowser.net/). For RNA-seq data, the ‘cpm’ algorithm of the “edgeR” package (V 2.12.0) was used to convert the count data into counts per million reads mapped (CPM), which was used to estimate the level of expression of each gene, before log2 transformations were performed. Based on the mutation data, the “Maftools” package (V 3.38.1) was used to analyze these results. In this study, we kept the expression profiles of the primary solid tumor samples, removed patients with missing survival information, and filtered out patients with less than 30 days of follow-up.

### Identification of Metabolic Subtypes

Glycolysis and cholesterol synthesis genes were obtained from Molecular Signatures Database v7.5 in the ‘REACTOME_GLYCOLYSIS’ (n = 72) and ‘REACTOME_CHOLESTEROL_ BIOSYNTHESIS’ (n = 25) gene set. After removing the genes with no expression in all samples, a total of 93 genes were obtained, including 69 glycolysis genes and 24 cholesterol biosynthesis genes. *ConsensusClusterPlus* V1.60.0 was used to cluster these genes. The parameters were as follows: reps = 100, pitem = 0.8, pfeature = 1, distance = “Spearman”, clusterAlg = “hc”, innerLinkage = “ward.D2”, finalLinkage = “ward.D2”. Median expression of z-scores of two co-expressed genes identified four metabolic subgroups. We defined them as quiescent (glycolysis ⩽ 0, cholesterogenesis ⩽ 0), glycolytic (glycolysis >0, cholesterogenesis ⩽ 0), cholesterogenic (glycolysis ⩽0, cholesterogenesis >0) and mixed (glycolysis >0, cholesterogenesis >0)[17]. To determine whether patients in these four types differed from each other, we performed principal component analysis (PCA). Survival analyses were then conducted, and log-rank test P values were also analyzed.

### Analysis of differential expression genes

Based on the survival analysis of the four subtypes, the two subtypes with the best and worst prognosis were selected for differential expression analysis using limma with threshold: FDR < 0.05 and log2| FC | > 1.5. GO, and Kyoto Encyclopedia of Genes and Genomes (KEGG) enrichment analysis was then performed on selected differentially expressed genes (DEGs), showing the results with FDR < 0.05.

### Prognostic gene selection based on machine learning

The TCGA-LUAD samples were randomly divided into training (80%) and test (20%) sets. And a chi-square test was performed on the training and test samples to show any significant difference between the two sets.

We applied DEGs to construct 5 Machine learning models, such as extreme gradient boosting (XGBoost), Random Forest Classifier (RFC), Logistic Regression (LR), Support Vector Machine (SVM), and K-Nearest Neighbors (KNN) were used to developed prediction models. Stratified k-fold cross-validation was used to validate the performance of the models.

In a predictive classification model, the most fundamental classification evaluation metric is based on comparing predicted and true values. For a binary classifying problem, comparing true and predicted classification results can categorize all samples into four classes: True Positive (TP), True Negative (TN), False Positive (FP), and False Negative (FN). To make the model perform better, a series of manually tuned real experiments were conducted. Various model training algorithms correspond to various hyperparameters, and the best hyperparameters were obtained by cross-validation in the training set.

We used global visualization of SHAP values to show the importance of the input features and selected the top nine most important features as the prognostic genes with the help of the best model.

### Construction and validation of a prognostic risk model

Using multivariate Cox regression analysis to construct a prognostic risk model based on the expression of prognostic genes. Then, we calculated the risk score for each sample as follows: 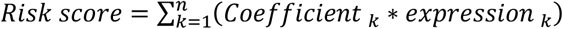. The samples were divided into high and low risk groups based on the median value of the risk scores. We validated the robustness of the model by using the training set and the test set and plot the risk score distribution as well as the ROC curve and KM survival curve. The relationship between risk scores and selected clinical characteristics was also analyzed by classifying high and low risk subgroups.

## 3. Results

### Identification of four metabolic subtypes of LUAD based on glycolysis and cholesterol synthesis gene expression

After screening, 501 LUAD tumor samples from TCGA were used to select co-expressed cholesterogenic and glycolytic genes (Figure 1A). When k=10, the genes in C3 were identified as glycolytic co-expression genes, all of which are in the glycolytic pathway (ALDOA, GAPDH, GCKR, GPI, PFKP, PKM, TPI1), and the genes in C10 were identified as cholesterogenic co-expression genes, all of which are in the cholesterogenic pathway (ACAT2, CYP51A1, HMGCR, HMGCS1, IDI1, MSMO1, SQLE). Samples were assigned to four subtypes (Figure 1B). PCA was performed to demonstrate differences between these subgroups (Figure 1C). Kaplan-Meier survival curve indicated significant prognostic differences among the four subtypes (log-rank test, P =0.0003) and the worst prognosis for patients in the mixed group and the best in the cholesterogenic group (Figure 1D). Figure 1E shows the co-expressed cholesterogenic and glycolytic genes expression of the four metabolic subtypes.

**Figure 1.**
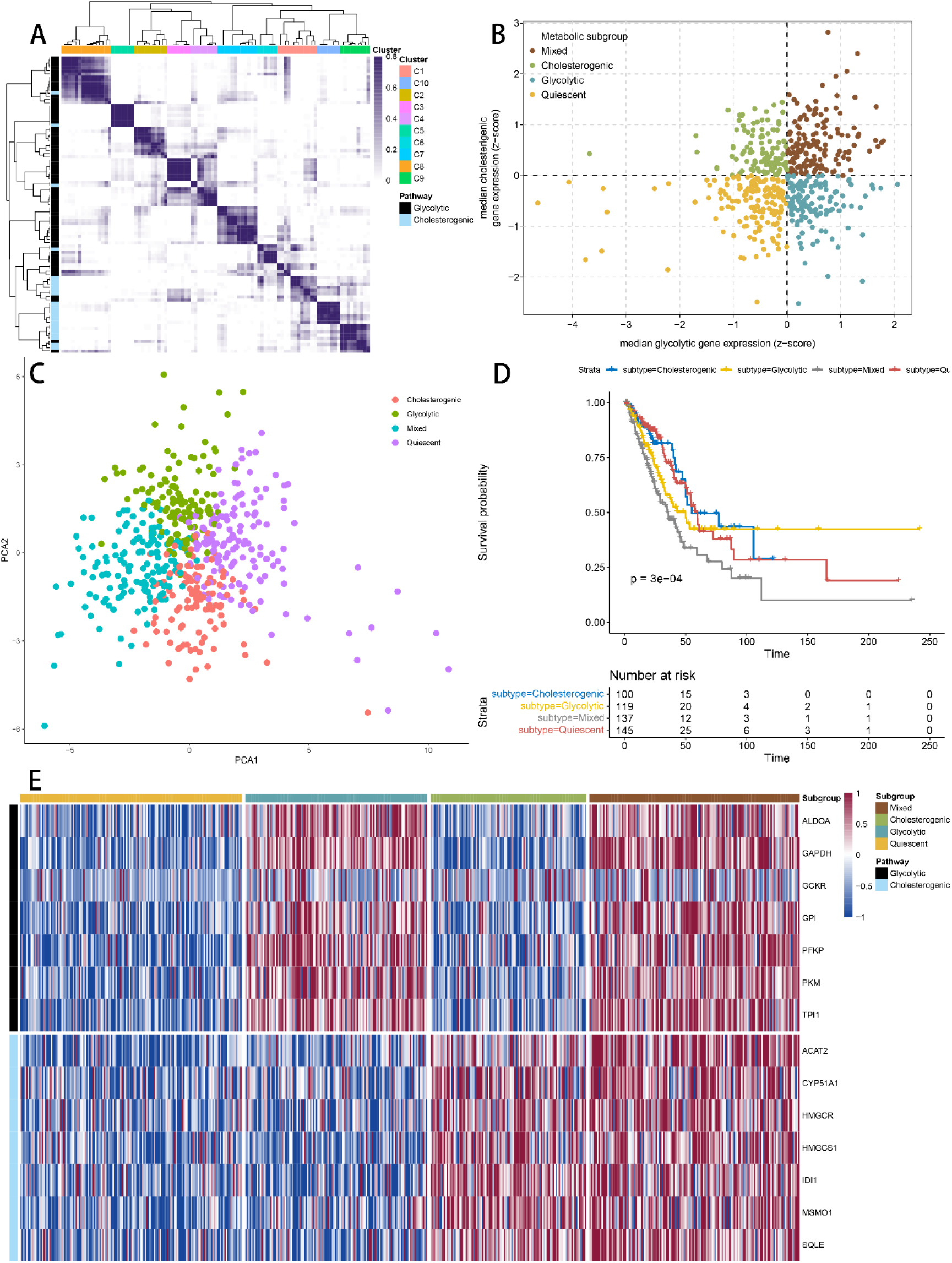
(A) Consistent clustering of glycolytic and cholesterogenic genes. (B) Sample classification according to median glycolytic and cholesterogenic genes expression. (C) Patients with different metabolic subtypes showed significant differences between each other by PCA. (D) Kaplan-Meier survival curves for different subgroups of patients. The clinical outcome endpoint was OS. (E) Heatmap of expression levels of co-expressed glycolytic and cholesterogenic genes in each metabolic subgroup. PCA, principal components analysis; OS, overall survival; P-value of <0.05 was considered statistically significant.

### Differential expression analysis between cholesterogenic and mixed groups

The cholesterol subtype had the best prognosis, in contrast to the mixed subtype, which had the worst in the four metabolic subtypes, suggesting that patients with high expression of only cholesterol-related genes had a better outcome, while those with high expression of glycolytic and cholesterol synthesis genes had a worse outcome. To further investigate the effect of cholesterol and resting subtypes on cancer prognosis, 445 DEGs were identified using the *“limma”* package (V3.52.2), of which 221 were up-regulated and 224 were down-regulated (Figure 2A). Figure 2B shows the heatmap of the 100 genes with the largest up- and down-regulation differences, respectively. GO enrichment analysis and KEGG pathway analysis were performed on 200 DEGs, and the annotated results (FDR<0.05) are visualized in Figure 2C-2D. The results showed that complement and coagulation cascades, cellular senescence, human T-cell leukemia virus 1 infection, p53 signaling pathway, microRNAs in cancer, and cell cycle are enriched significantly.

**Figure 2.**
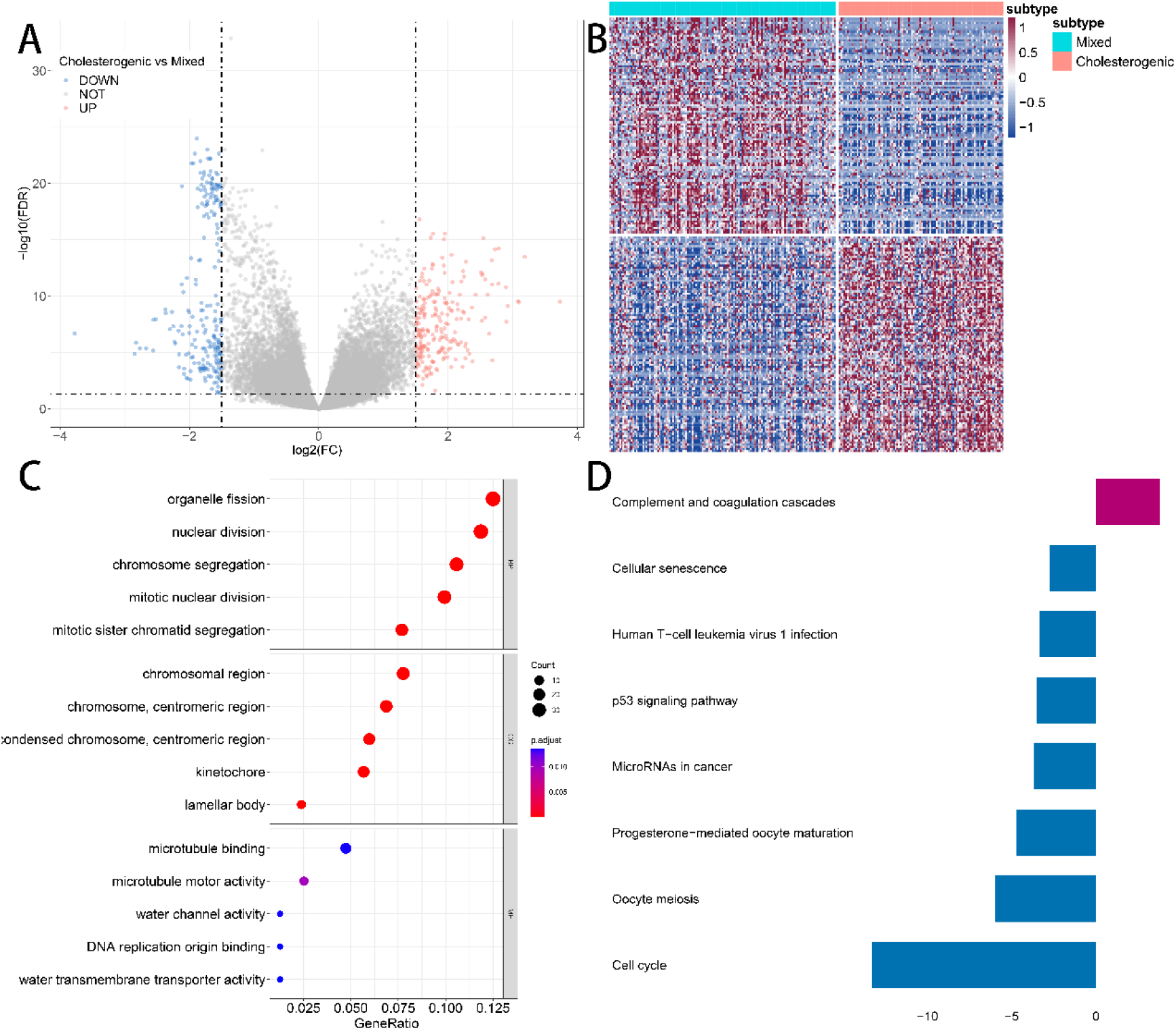
(A) Volcano map of differentially expressed genes in the TCGA dataset for cholesterogenic and mixed subtypes. (B) Heatmap of differentially expressed genes in the TCGA dataset for cholesterogenic and mixed subtypes. (C) Annotation using GO for differentially expressed genes. (D) Annotation using KEGG for differentially expressed genes.

### Prognostic gene selection based on machine learning

a total of 501 participants were analyzed, among which 320 were alive; these 501 participants were randomly split into training and test datasets. The results of all Machine learning models for predicting survival status are shown in Figure 3A. The XGBoost model had the better performance in the test set.

**Figure 3.**
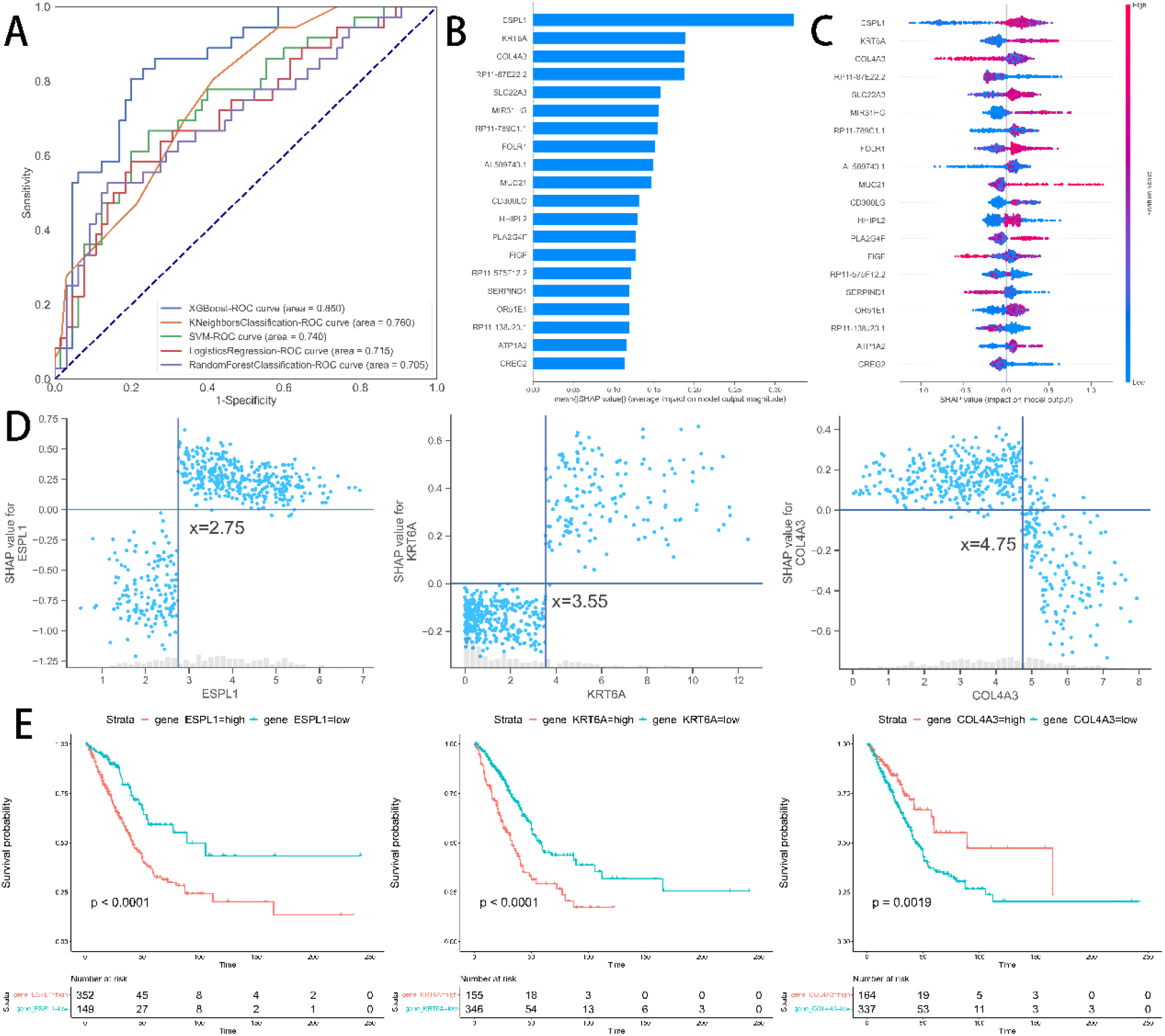
(A) Receiver operating characteristic curves showing the performance of the five models. (B) Histogram of mean Shapley additivity explained (SHAP) value for the top 20 predictors. (C) bee-swarm plots. The X-axis is the SHAP value, representing the impact of the outcome predictors on the Y-axis. One dot represents one individual patient. The higher the SHAP value, the poorer prognosis. (D) The dependence plot of the ESPL1, KRT6A, and COL4A3. (E) Three gene KM curves on the training set.

In Figure 3B-3C, the summary plot of the mean absolute SHAP values shows the order of importance of the features from highest to lowest with the help of the XGBoost model. It shows that ESPL1, KRT6A, COL4A3, RP11-87E22.2, SLC22A3, MIR31HG, RP11-789C1.1, FOLR1, and AL589743.1 are the nine most important genes for predicting survival status. Figure 3D shows the association between the SHAP values of the three most important genes and the expression of each gene. The truncation thresholds for these three predictor variables can be defined from the graph to distinguish between good prognosis (i.e., SHAP-value < 0) and poor prognosis (i.e., SHAP-value > 0). ESPL1 beyond 2.75 or KRT6A greater than 3.55 or COL4A3 less than 4.75 increases the SHAP value and thus the poorer prognosis. Kaplan-Meier survival curves were plotted for high and low risk groups according to the gene cutoff values, and the results indicate significant prognostic differences between the groups determined by the SHAP values (Figure 3E).

### Model construction and evaluation

Using a multivariate cox regression model the nine-gene signature equation is as shown below: risk score**=** 1.32*ESPL1 + 1.11*KRT6A + 0.96*COL4A3 + 0.7*RP11-87E22.2 + 1.10*SLC22A3 + 1.22*MIR31HG + 1.08*RP11-789C1.1 + 0.96* AL589743.1 +1.06*FOLR1. Risk scores were calculated using prognostic gene expression levels for each sample, and risk score distributions and gene expression heatmap of the training set were plotted (Figure 4A). Survival times were significantly shorter in the high-risk score samples than in the low-risk score samples, suggesting that samples with higher risk scores are more likely to have a poor prognosis. Two hundred samples with risk scores greater than the median were classified as a high-risk group, and 200 samples less than the median were classified as a low-risk group. Significant prognostic differences existed between the high- and low-risk groups (p<0.05, Figure 4B). Figure 4C shows the predicted classification performance based on risk scores for 1, 3, and 5 years.

**Figure 4.**
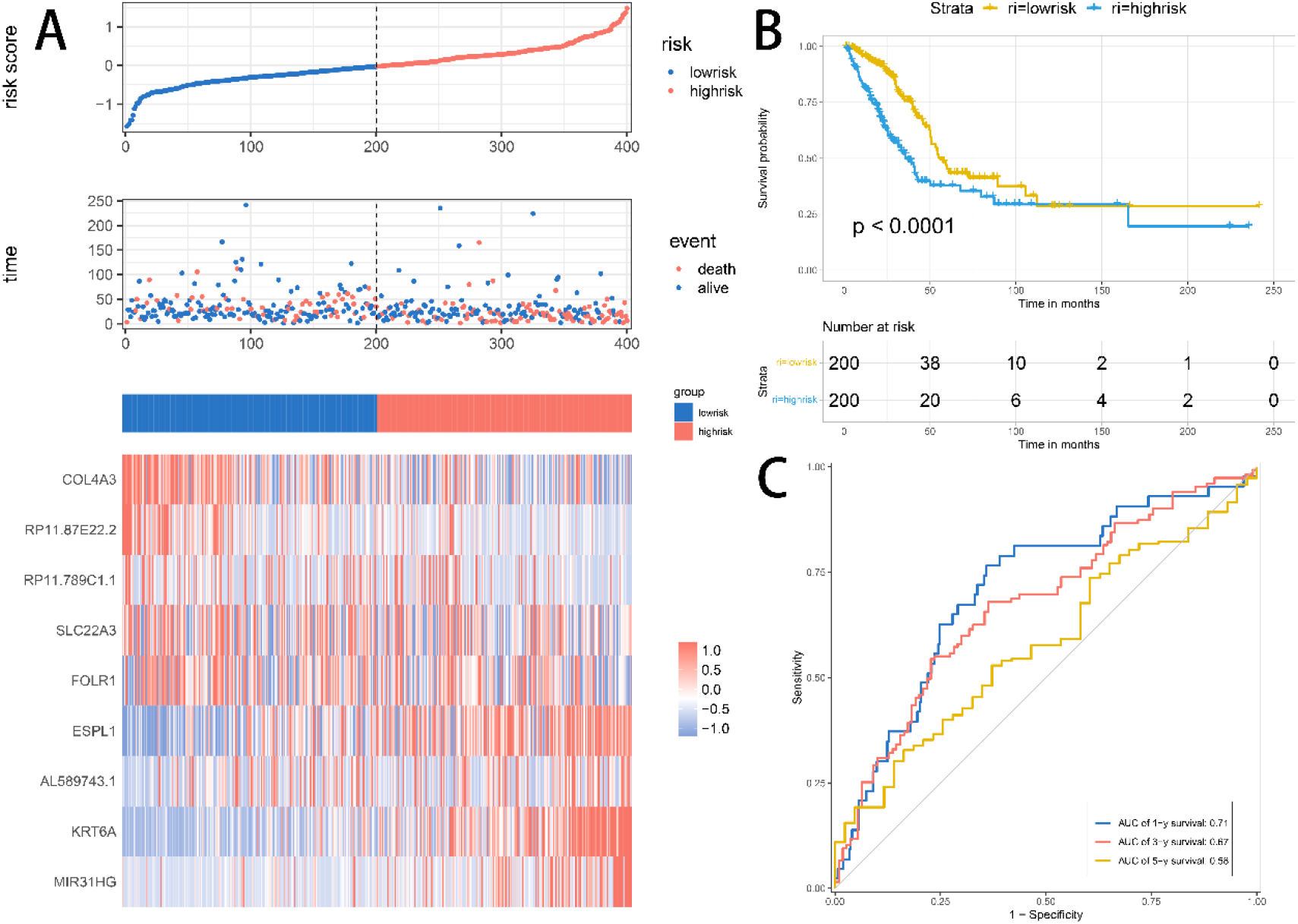
(A) Distribution of risk scores, survival time, and heatmap of expression of 9 genes in the training set. (B) The KM survival curve of the risk model in the training set. (C) The ROC curve of the risk model in the training set.

Risk score distributions and gene expression heatmap of the test set were shown in Figure 5A. Consistent with the trend in the training set, indicating a better prognosis for samples with low risk scores. The KM survival curve demonstrated a significant prognostic difference between the high-risk and low-risk groups in the test set (Figure 5B). Figure 5C shows the predictive classification performance based on risk scores at 1, 3, and 5 years.

**Figure 5.**
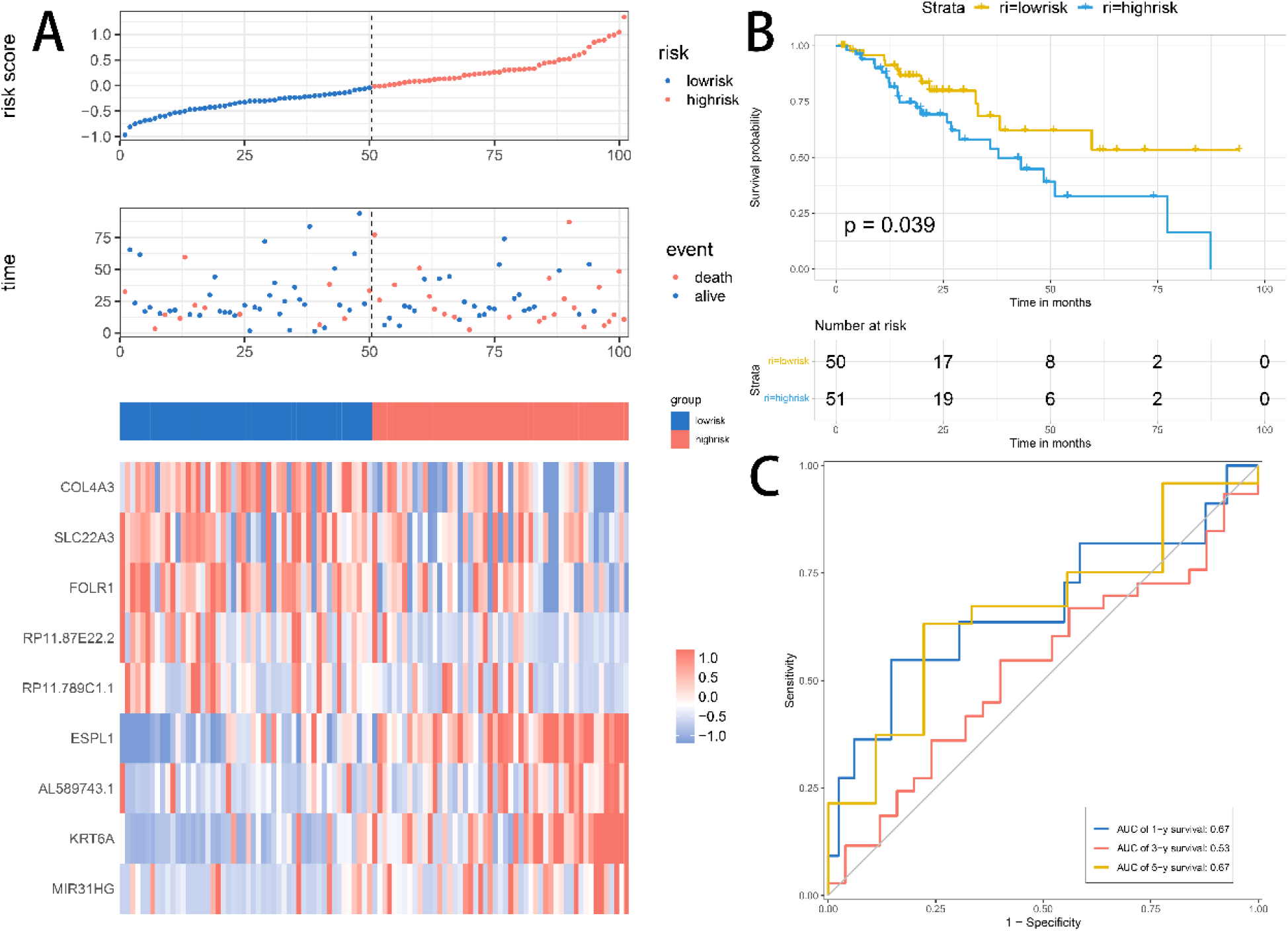
(A) Distribution of risk scores, survival time, and heatmap of expression of 9 genes in the test set. (B) The KM survival curve of the risk model in the test set. (C) The ROC curve of the risk model in the test set.

### Relevance analysis of risk models to clinical characteristics

Risk scores were associated with clinical characteristics significantly (Figure 6A). By gender, risk scores were significantly higher in the male group than in the female group; by stage, the higher the stage, the higher the risk score; by subtype, risk scores were higher for the mixed subtype with a poor prognosis and lower for the cholesterol subtype with a good prognosis. Therefore, the prognostic model has shown good predictive performance with respect to different clinical characteristics. According to the risk score, age can be classified into high and low risk groups with significant prognostic differences (p<0.05, Figure 6B). A waterfall plot was used to present mutation data for each gene in high and low risk groupings per sample (Figure 6C).

**Figure 6.**
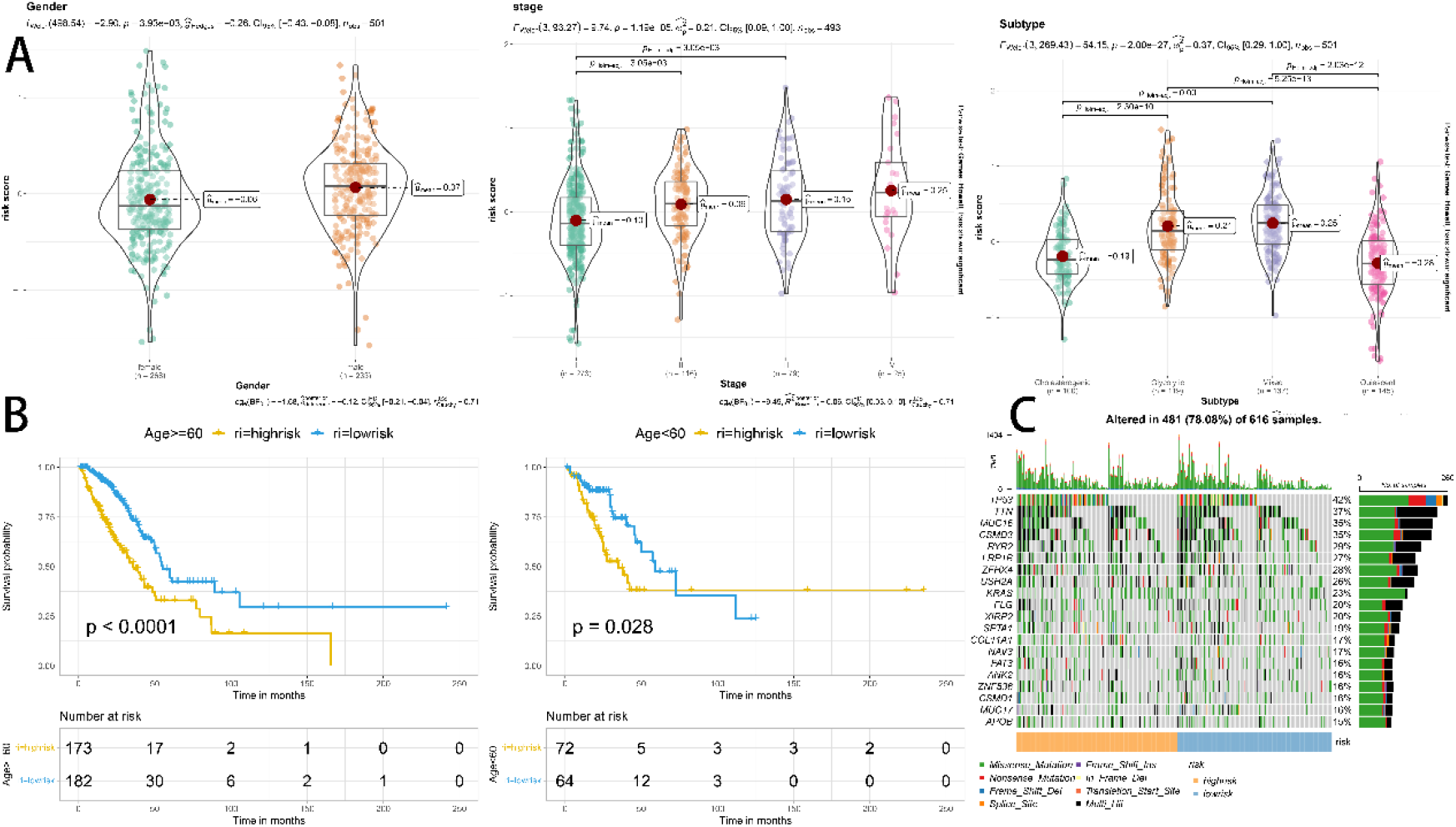
(A) Association of risk scores between samples of various gender, stage, and subtype groups. (B) Prognosis comparison of various age groups according to risk scores. (C) Waterfall plot displaying details of mutations in each gene for each sample in the high and low risk groups.

## 4. Discussion

Lung cancer is one of the most deadly malignancies in humans, and most patients with advanced lung cancer experience recurrence and treatment resistance. The abnormal metabolism of cancer cells characterized by high glycolysis that occurs even in the presence of high amounts of oxygen, a metabolic reprogramming called the Warburg effect or aerobic glycolysis, has been recognized as a new hallmark of cancer [18]. Inhibition of glycolysis is considered a therapeutic option for aggressive cancers, including lung cancer, and related genes can be used as potential targets for metabolic therapy against cancer cells, such as ARID1A and circ-ENO1[19–21]. Altered metabolism is not limited to cellular energy pathways but also includes alterations in lipid biosynthesis and other pathways (e.g., polyamine processing) in lung cancer and can affect its surrounding microenvironment[22]. It has been shown that lung cancer tissues demonstrate elevated cholesterol levels because the proliferation of cancer cells depends heavily on its availability. Strategies to reduce cholesterol synthesis or inhibit cholesterol uptake have been proposed as potential antineoplastic therapies[23,24]. Therefore, it is essential to clarify the metabolic pathways of lung cancer for its prevention and treatment.

In this study, based on 93 glycolysis and cholesterol synthesis genes, in order to find the most representative genes, consistent clustering was used to minimize the gene numbers, yielding 7 and 7 cholesterogenesis and glycolysis co-expressed genes, respectively. Based on these genes, the samples were classified into four subtypes: glycolytic, cholesterogenic, quiescent, and mixed. Although cholesterol plays a crucial role in tumors, survival analysis showed that cholesterol subtypes have a better prognosis than other subtypes, and randomized controlled trials could not support a survival benefit through lipid lowering in lung cancer patients, the reasons for which deserve further investigation [25].

We identified DEGs between the best and worst prognosis subtypes and performed a functional enrichment analysis. The results showed significant enrichment of DEGs between the mixed and cholesterogenic subtypes in terms of p53 signaling pathways, microRNAs in cancer, and cell cycle. Then we decided to build the model using machine learning, with DEGs as features and ending events as labels. It performs best in the test set based on XGBoost’s powerful ability to handle complex classification problems. SHAP was then used to select the most important nine features. Then the most important nine features which SHAP selected were used to construct a prognostic model using a multivariate cox regression model. And this makes it possible to combine the excellent classification power of machine learning with the interpretability of the prognostic model.

Nine prognostic genes were included four non-coding RNA genes (RP11.87E22.2, RP11.789C1.1, AL589743.1, and MIR31HG) and five coding protein genes (COL4A3, SLC22A3, FOLR1, ESPL1, and KRT6A). After reviewing the kinds of literature, we found that the role of many prognostic genes has been studied concerning lung cancer and has been revealed to impact tumorigenesis and progression. Specifically, ESPL1 expression was positively correlated with SHAP values, and high expression of ESPL1 has been previously shown to be associated with poor prognosis in lung cancer by Zhao et al. [26]. Similarly, KRT6A, MIR31HG, and FOLR1 have been found to enhance lung cancer proliferation and may be potential therapeutic targets[27–29]. Subsequently, all samples were classified into high and low risk groups, and the clinical characteristics of the different risk groups were analyzed. Consistent results with the training set were observed in this test set. The model was robust on the training and test datasets and had a great predictive performance.

There are also some limitations to this study. Firstly, the performance of our model has not been tested externally, and there are doubts about its availability for large-scale use. Secondly, the biological associations between the selected prognostic genes remain to be investigated, and their biological explanations with prognostic profiles are to be explored. Future experimental verification is needed. Finally, using the median risk score as a cutoff value to classify high and low risk needs to be optimized.

## 5. Conclusions

Patients with LUAD were effectively typed by glycolytic and cholesterogenic genes and were identified as having the worst prognosis in the glycolytic and cholesterogenic enriched gene groups. Prognostic genes selected by the XGboost algorithm and SHAP analysis can be used to analyze patient prognosis. The prognostic models can provide an important basis for clinicians to predict clinical outcomes for patients.

